# The fidelity of transcription in human cells

**DOI:** 10.1101/2022.05.10.491385

**Authors:** C. Chung, B.M. Verheijen, X. Zhang, B. Huang, A. Coakley, E. McGann, E. Wade, O. Dinep-Schneider, J. LaGosh, M. Anagnostou, S. Simpson, K. Thomas, J. M. Ernst, A. Rattray, M. Lynch, M. Kashlev, B. A. Benayoun, Z. Li, J. Strathern, J-F. Gout, M. Vermulst

## Abstract

Accurate transcription is required for the faithful expression of genetic information. To provide insight into the molecular mechanisms that control the fidelity of transcription, we analyzed the landscape of transcription errors in human embryonic stem cells. These measurements provide the first reasonable estimate of the fidelity of transcription in human cells and identify multiple genetic and epigenetic factors that control its accuracy. In addition, we developed a new reporter mouse to identify cell types and tissues that commit these errors the most. These experiments revealed that CA1 and dentate gyrus neurons are highly sensitive to transcriptional mutagenesis, lending new support to the hypothesis that transcription errors play a role in the progression of Alzheimer’s disease. Taken together, these experiments provide unprecedented insight into the fidelity of gene expression in human cells and the molecular mechanisms that govern the central dogma of life.

## INTRODUCTION

The human genome provides a precise, biological blueprint of life. To implement this blueprint correctly, it is essential that our genome is transcribed with the utmost precision. Sporadic errors are unavoidable though, and these errors reveal how important transcriptional fidelity is for cellular health. For example, in patients with non-familial cases of Alzheimer’s disease, transcription errors generate toxic APP and UBB peptides that are part of the amyloid plaques that characterize the disease^1,2^, suggesting that they contribute to disease progression. The toxic UBB peptide (UBB^+1^) is also a potent inhibitor of the proteasome ubiquitin complex^3^ and can be found in protein aggregates from patients with Guam parkinsonism–dementia complex, tauopathies and polyglutamine repeat disorders^1,4-6^, suggesting that it contributes to other protein misfolding diseases as well. Intriguingly, these errors occur repeatedly at the same location, creating identical mutant proteins that can partially mimic the impact of a mutation on cellular health. Random transcription errors (errors that occur only once at each position) contribute to protein misfolding diseases as well. While studying yeast cells that display error-prone transcription, we found that these errors tend to compromise the structural integrity of proteins and induce protein misfolding at a global scale^7^. Although most of these misfolded proteins are benign, their sheer volume can overwhelm the protein quality control machinery and prevent the degradation of toxic proteins that are normally targets for this machinery, including Aβ1-42 (AD), TDP-43 (amyotrophic lateral sclerosis and frontotemporal dementia, ALS, FTD), HTT103Q (Huntington’s disease) and RNQ1^7^ (a prion).

Taken together, these observations suggest that transcription errors can affect proteotoxic diseases by two complementary mechanisms. They can generate specific peptides that are associated with the disease, and they can create the environment that allows these peptides to persist and seed aggregates. Transcription errors affect other cellular processes as well though. For example, they can induce oncogenic pathways in human cells^8^, limit the lifespan of yeast^7^, compromise the metabolism of NAD, nucleotides and amino acids^9^, and change the fate of bacterial cells^10^. However, despite our increased understanding of the impact of transcription errors on cellular health, relatively little is known about the molecular mechanisms that control the fidelity of transcription, especially in human cells. To address this issue we measured the error rate of transcription in H1 human embryonic stem cells (H1 hESCs). These experiments reveal various genetic and epigenetic factors that modulate the fidelity of transcription in human cells, demonstrate that the fidelity and strength of gene expression are linked, and suggest that the DNA repair protein BRCA1 has a previously uncharacterized role in the suppression of transcriptional mutagenesis. In addition, we developed a new reporter mouse that revealed that neurons implicated in Alzheimer’s disease are prone to transcriptional mutagenesis, strongly supporting a 20-year-old hypothesis that suggests that transcription errors play an important role in the etiology of this disease^1,2^.

## RESULTS

### Data overview

To determine how accurately the human genome is transcribed, we sequenced the transcriptome of H1 hESCs with an optimized version^9^ of the “circle-sequencing assay”^11,12^ (**fig. 1A**). We then compared their transcriptome to a custom-made reference genome (300x coverage) to identify transcription errors (**fig. 1B-C, S1**). In addition, we used these datasets to demonstrate that all replicates had a similar transcriptomic profile, indicating that their health and overall status were comparable (**fig. S2)**. This analysis yielded 101,884 transcription errors, distributed over every major RNA species in human cells (**fig. 1C**), including mRNA, rRNA, tRNA, small nuclear RNA, small nucleolar RNA (snoRNA), ribozymes and various classes of pseudogenes. Because these transcripts are synthesized by different RNA polymerases (RNAPs), this dataset allowed us to determine the accuracy of every major nuclear RNAP in human cells (**fig. 1E-G, 2A**). In addition, we detected a substantial number of transcripts that were derived from the mitochondrial genome (mtDNA), which allowed us to determine the accuracy of the mitochondrial RNAP as well (**fig. 1E-G, 2A**). Finally, we combined these errors into a single file, so that they can be downloaded through an internet link and visualized as a track in the UCSC genome browser.

**Figure 1.**
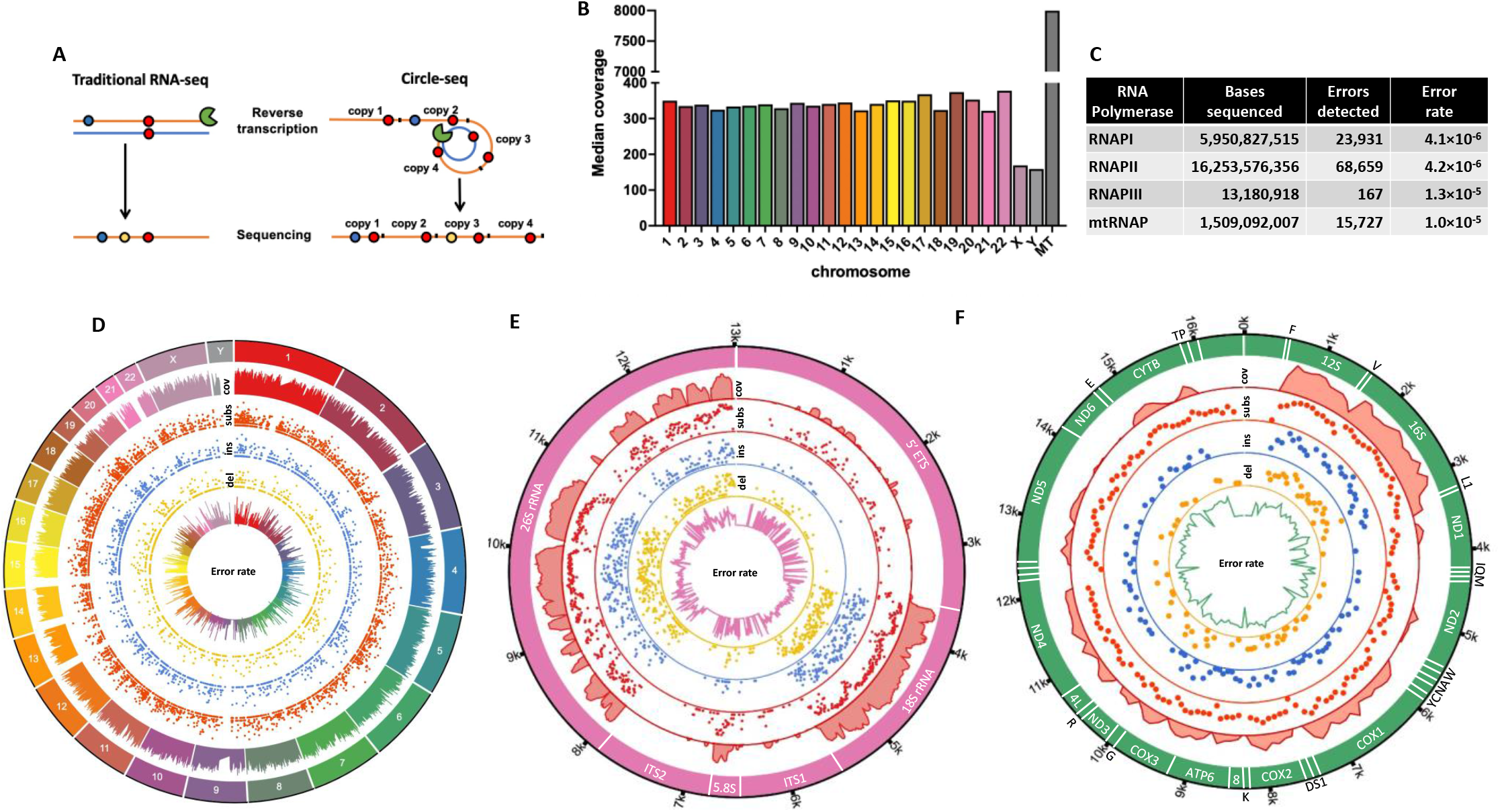
Overview of dataset. **A**. Concept of the circle-sequencing assay. Traditional RNA-seq assays introduce reverse transcription errors (blue circles) during the generation of cDNA libraries (orange lines, the RNA template is in blue) that are indistinguishable from true transcription errors (red circles). In addition, sequencing mistakes (yellow circles) arise during massively parallel sequencing reactions that can be misinterpreted for transcription errors. To circumvent these pitfalls, RNA molecules are ligated to themselves, and circularized prior to reverse transcription. These circular molecules can then be reverse transcribed in a rolling circle fashion to generate linear cDNA molecules that consist of multiple concatenated copies of the original RNA template. If a transcription error was present in the template, this error will be present in every concatenated copy, while reverse transcription errors or sequencing mistakes will only be present in one or two copies so that true transcription errors to be distinguished from artifacts. **B**. The median coverage per base of each chromosome by DNA-sequencing to generate a reference genome. **C**. The distribution of RNA bases sequenced and the errors detected in transcripts that were synthesized by different RNA polymerases. **D**. A heatmap of all transcripts detected in each of the 6 replicates. **E-G**. Transcription errors detected in mRNAs (**E**), rRNA (**F**) and mitochondrial RNA (**G**). Genes and chromosomes are located on the outer ring, followed by the coverage of these sequences, base substitutions per 10kb or 100bps in red, insertions per 10kb or 100bps in blue, deletions per 10kb or 100bps and the error rate/bp in yellow over these intervals. The inner ring represents the error rate.

### Human cells prioritize the fidelity of specific transcripts

To determine the parameters that control the error rate of transcription, we first compared the accuracy of different RNA polymerases to each other. This analysis demonstrated that RNAPI (4.1×10^−6^/bp) and II (4.2×10^−6^/bp) make the least mistakes, followed by the mitochondrial RNA polymerase (mtRNAP, 1.0×10^−5^/bp) and RNAPIII (1.3 ×10^−5^/bp). Thus, human cells prioritize the fidelity of RNAPI and II over other polymerases. We previously observed a similar trend in other organisms, indicating that this feature is evolutionarily conserved^9^. The increased error rates of RNAPIII and mtRNAP are fueled by the unique error spectra of these polymerase. For example, RNAPIII makes 5 times more G→A errors than RNAPI or II, which drives their error prone phenotype (**Fig. 2B-D**). It is important to note though, that despite the relatively high fidelity of RNAPI and II, the error rate of transcription is still >100-fold higher than the mutation rate^13^, underscoring the vast potential of transcription errors to create mutant proteins.

**Figure 2.**
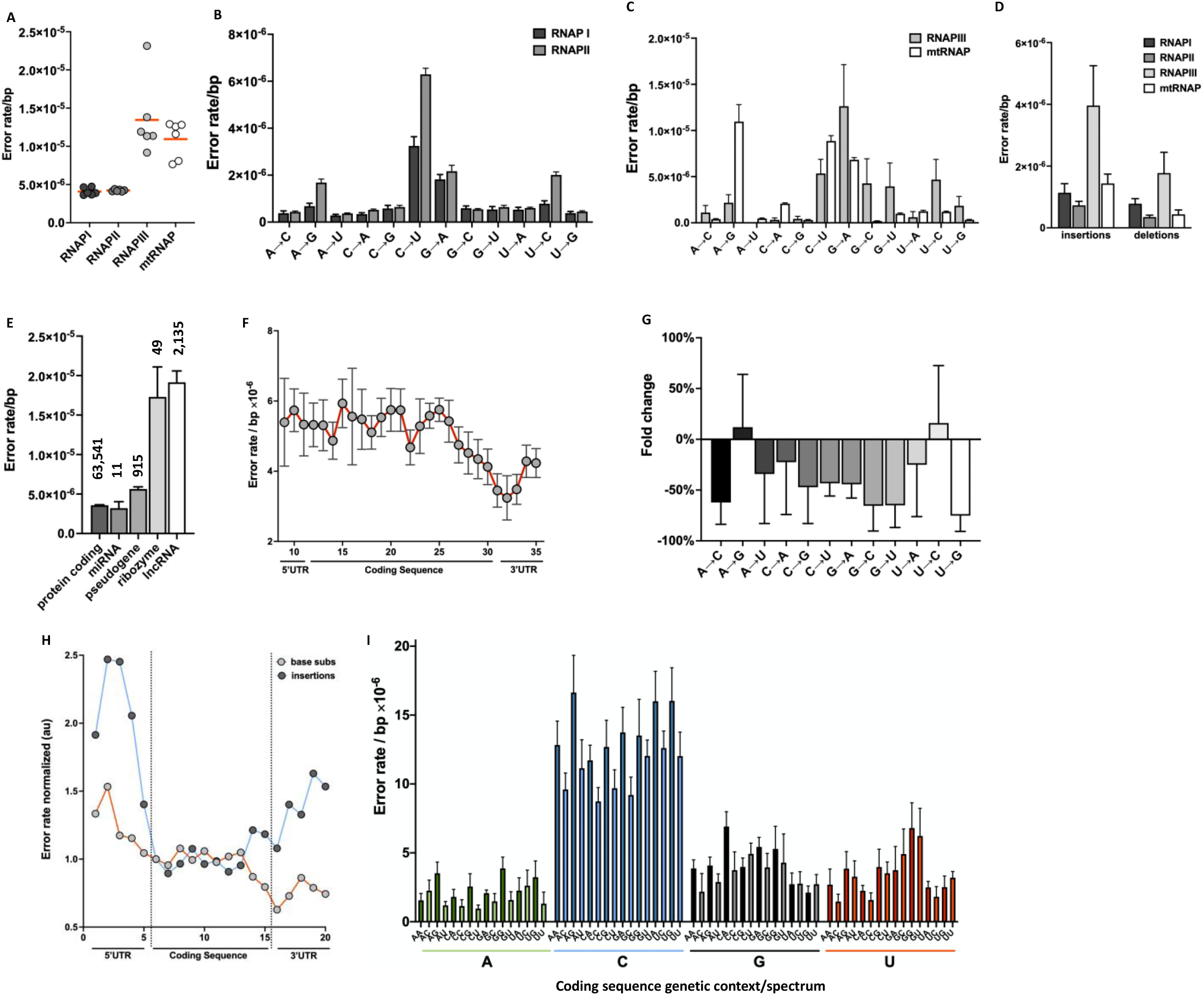
Genetic determinants of the error rate of transcription. **A-D**. Different RNA polymerases display different error rates (**A**) and spectra (**B-D**). **E**. Different transcripts generated by RNAPII display different error rates. The numbers above the bars indicate the number of errors detected in each class of RNAs. **F**. The error rate of transcription decreases near the stop codon. **G**. Most single base substitutions decrease near the stop codon (interval 25-32 in **2F**). **H**. Insertions increase near the stop codon. Please note that the insertion and base substitution error rate have been normalized to each other, to better display the qualitative differences between them. In addition, the intervals have been condensed by a factor of 2 compared to **2F**. For example, interval 10 equals interval 20 in figure 2F. **I**. The error rate of transcription is highest if the next base that needs to be inserted is a pyrimidine.

Since our data suggests that human cells prioritize the fidelity of some transcripts over others, we wondered how expansive this prioritization process is. To explore this concept, we examined how accurate RNAPII is when it transcribes different classes of genes. This analysis revealed that RNAPII makes less mistakes when it transcribes genes that encode miRNAs and proteins compared to pseudogenes, ribozymes and lncRNAs (**Fig. 2E**). To gain more insight into the molecular mechanisms that might be responsible for this observation, we analyzed the error spectrum of RNAPII on ribozymes and lncRNAs, the two templates with the highest error rates. Interestingly, we found that transcripts that encode ribozymes display a sharp increase in the rate of insertions, deletions, and G→A errors, a signature that is highly reminiscent of RNAs transcribed by RNAPII without its fidelity factors^9^ (**fig. S4 A-B**). Thus, one potential mechanism by which RNAPII could alternate the fidelity of transcription is the inclusion or exclusion of fidelity factors.

A different mechanism seems to be responsible for the increased error rate of lncRNAs, because in contrast to ribozymes, these RNAs display an increased rate for every possible transcription error (**fig. S4 C-D**). Intriguingly, we made the inverse observation when we examined the fidelity of transcription in protein coding genes. Surprisingly, we found that the error rate of transcription is not equal across the length of these genes. Instead, nearly every error decreases in frequency as RNAPII approaches the stop codon (**Fig. 2F-G**). Because so many errors fluctuate at these locations, it is unlikely that these observations are caused by an external source of transcription errors. For example, DNA damage is a powerful source of transcription errors^14^. However, an increase or decrease in DNA damage is expected to cause only one, or a few errors to change in frequency^15^. One potential mechanism to alter the error rate of many, or all possible errors, would be to alternate the speed of RNA synthesis. The speed and accuracy of RNA^16,17^ and DNA polymerases^18^ are inversely correlated, so that the faster a polymerase works, the less accurate it is. This hypothesis is examined further in the sections below. It is important to note though, that in contrast to base substitutions insertions and deletions do increase in frequency near the stop codon (**Fig. 2H, fig. S5**). Most likely, this observation is caused by a reduction in the efficiency of nonsense-mediated mRNA decay (NMD). An important distinction between indels and base substitutions is that indels frequently result in premature stop codons that trigger for NMD^19^. However, it is increasingly difficult for a cell to distinguish between a premature stop codon and a native stop codon if these codons are located close to each other. Accordingly, the increased number of insertions and deletions near the native stop codon is most likely due to the reduced ability of NMD to detect these events. We previously confirmed this idea in yeast cells^9^.

### The primary sequence of the human genome alters the fidelity of transcription

Next, we wondered whether the fidelity of transcription is affected by the primary sequence of the human genome. To answer this question, we examined the error rate of transcription in different genetic contexts. First, we computed the error rate of RNAPII on all 4 nucleotides, and then tested whether these error rates alter as a result of their 5’ and 3’ neighbors. This analysis indicated that that was indeed the case and that the 3’ base is the most important determinant for the fidelity of transcription (**fig. 2I**). If the next base that needs to be inserted is a pyrimidine, the error rate tends to be lower than when it is a purine. Because the incorporation of a pyrimidine is slightly slower than a purine, it is possible that the speed of incorporation plays a role in this phenomenon as well^20^. A slower incorporation rate would provide RNAPII with more time to remove a misincorporated base. Regardless, we observed a similar trend in yeast cells, indicating that this molecular phenomenon is evolutionarily conserved^9^. Finally, we created similar genetic context maps for rRNA and mtRNA **(fig. S6)**.

We also examined the error rate of transcription on tracts of mono- and di-nucleotide repeats. RNAPII tends to slip on these tracts in model organisms^9,21,22^ and similar slippage events have been observed in humans. For example, in patients with Down syndrome and non-familial cases of Alzheimer’s disease, RNAPII was found to slip on two dinucleotide tracts in the *APP* and *UBB* gene, resulting in shortened peptides that are part of the amyloid plaques that characterize the disease^1,2^. However, it is unknown how frequently these slippage events occur in human cells, which obscures their impact on human aging and disease. To fill this gap in our knowledge, we measured the error rate of RNAPII on mono and di-nucleotide repeats and compared it to the rest of the human genome. This analysis demonstrates that repeats raise the error rate of insertions and deletions up to a 200-fold (**Fig. 3A-C**), an increase that is proportional to the length of the tracts. In addition, we found that some repeats result in smaller, but significant increases in base substitutions (**Fig. 3A-C**). These observations suggest that toxic APP and UBB peptides are generated at an accelerated rate in human cells, which could promote the development of proteotoxic diseases.

**Figure 3.**
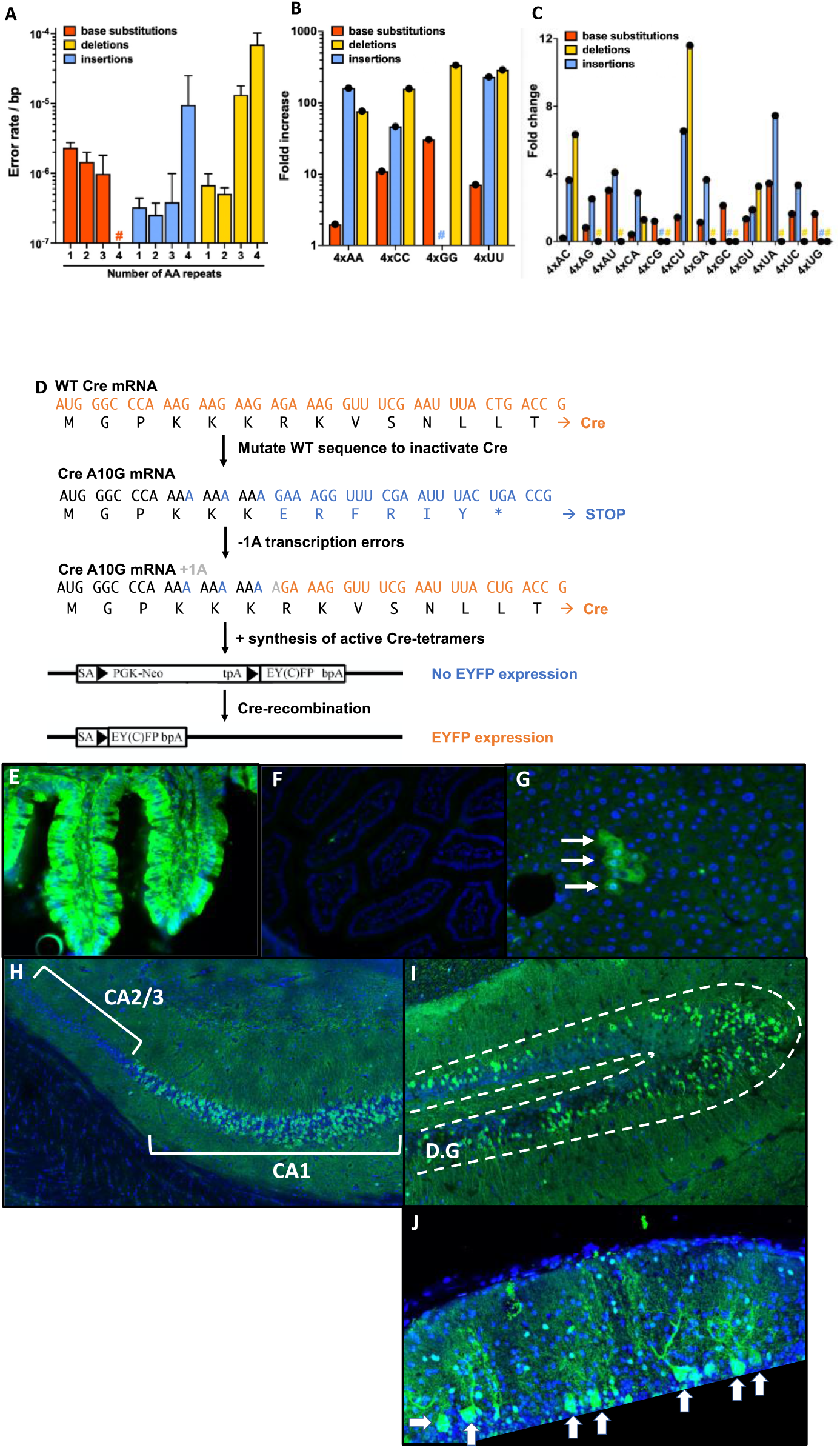
RNA polymerase II slips on mono and di-nucleotide tracts. **A**. Increased rates of insertions and deletions along a mono-nucleotide tract of adenines. Slippage events increase on sequences that are made up of 6 or 8 thymine bases in the DNA. **B**. Increased transcription errors on all possible mono-nucleotide tracts made up of 8 bases. **C**. Increased transcription errors on di-nucleotide tracts made up of 8 bases. Please note that for panel **B** and **C**, the WT and mutant bases of all 6 replicates have been combined together because of the relatively low coverage of these repeat sequences across the transcriptome. **D**. Genetic construct inserted in C57Bl/6j mice that contains a mutated version of the Cre-gene. This gene has been placed out of frame and contains a slippery tract of 10 adenines in a row. Upon an insertion or deletion that places Cre back in frame, a short burst of WT Cre-proteins is generated that can excise a Neo-cassette that interrupts an EYFP gene, leading to constitutive EYFP expression. **E**. A positive control where EYFP was activated in every intestinal cell by a Sox2 driven WT-Cre. **F**. EYFP expression is nearly absent in mice that express the Cre^out^ gene. **G**. EYFP cells are also sparse in the liver. Here, s single patch a EYFP liver cells is depicted, most likely representing a single transcription error that activated EYFP in one cell, which then clonally expanded. **H-I**. Large numbers of EYFP^+^ cells in the CA1 but not CA2 or CA3 region of the hippocampus (**H**), the dentate gyrus (**I**, D.G.) and cerebellum (**I**, Purkinje cells) of Cre^out^; EFYP^neo^ mice.

### Cells that are implicated in Alzheimer’s disease are prone to transcriptional mutagenesis

Because slippage events are implicated in Alzheimer’s disease and other protein misfolding diseases, we decided to investigate which cell-types commit these errors the most. To do so, we developed a new mouse model that can be used to identify cell types and tissues that have experienced slippage events. This mouse model carries a *Cre*-recombinase gene that was placed out of frame by a slippery tract of 10 adenines (Cre^out^, **Fig. 3D**). Because of this frameshift, the *Cre* gene can only produce functional Cre proteins if RNAPII slips on the adenine tract and places the *Cre* transcript back in frame. The proteins made from this transcript can then form a biologically active tetramer and excise a cassette elsewhere in the genome that permanently activates EYFP (EYFP^neo^). Thus, the end result of these transactions is that a single, transient transcription error is converted into an easily tractable genetic mutation. Note that translation errors cannot result in EYFP activation because they only generate a single Cre protein, which is insufficient to create a biologically active tetramer. And even though mutations could result in EYFP activation, these events occur 100-fold less frequently than transcription errors, which makes it unlikely that they result in large amounts of EYFP^+^ cells. For example, direct sequencing of EYFP^+^ neutrophils demonstrate that they do not carry mutations in the Cre gene, indicating that EYFP activity is not the result of genetic mutations (**J.S. unpubl. results**). We then aged Cre^out^; EYFP^neo^ mice for 1 year to allow EYFP^+^ cells to accumulate and stained intestine, liver and brain tissue with EYFP-antibodies (**Fig. 3E-I**). Interestingly, we found that liver, intestine and most brain regions contained few EYFP^+^ cells (**Fig. 3F-G**). However, EYFP^+^ cells did accumulate in specific structures of the brain. Most notably, we detected many EFYP^+^ cells in the dentate gyrus and CA1 neurons of the hippocampus (**fig. 3H-I**), brain regions that are pivotal to memory formation and the pathology of Alzheimer’s disease^23^. In contrast, neighboring CA2/3 cells displayed significantly less EYFP^+^ cells, indicating that CA1 and dentate gyrus neurons are uniquely error prone. These results indicate that in addition to the genes involved in Alzheimer’s disease, the affected cells are error prone as well. Together, these error prone phenomena could create a constant stream of toxic APP and UBB peptides that contribute to Alzheimer’s disease progression. In addition, we identified numerous EYFP^+^ Purkinje cells in the cerebellum (**fig. 5d**). Importantly, Purkinje cells are highly sensitive to protein aggregation as well, and a primary target for prion disease^24,25^, suggesting that error prone transcription could be a feature of multiple cell types that are sensitive to proteotoxic stress.

### The strength of gene expression is directly coupled to its fidelity

Next, we wondered how epigenetic factors might affect the fidelity of transcription as well. To answer this question, we cross-referenced the errors we detected with epigenetic modifications that were catalogued in H1 ESCs by the human ENCODE project^26^ and the NIH Roadmap Epigenomics Mapping Consortium. This analysis revealed that several histone modifications associated with active transcription, including H3K4me1, H3K4me2 and H3K4me3 marks, correlate with increased rates of base substitutions (**fig. 4A)**, insertions and deletions **(fig. S7**). To determine if the opposite is true as well (that markers associated with gene repression correlate with reduced error rates), we quantified the relationship between transcription errors and two mutually exclusive modifications on lysine 9 of histone 3 (H3K9). At this location, histones can either be acetylated (H3K9ac), which is a mark associated with active transcription, or tri-methylated (H3K9me3), which is a mark associated with repressed transcription^27^. We again found that the active H3K9ac mark was associated with an increased error rate, but in addition, we discovered that the repressive H3K9me3 mark was associated with a reduced error rate (**fig. 4A**). Taken together, these observations suggest that the strength and fidelity of gene expression are directly coupled to each other. One way to couple the fidelity of transcription to the strength of gene expression is the elongation speed of RNAPII: the faster an RNA polymerase works, the less precise it is^16^. To explore this hypothesis further, we examined the error rate near two histone marks that are associated with increased speed of elongation (H3K79me2 and H4K20me1^28^). Consistent with a role for elongation speed in transcriptional fidelity, we found that regions that carry these marks are transcribed less accurately than regions that do not (**fig. 4A**). Moreover, H3K79me2 and H4K20me1 marks tend to be lost near the 3’ end of genes^27^, locations where elongation is known to slow down due to a pile-up of RNAPs that are in the process of transcription termination^29^. Accordingly, these observations also support the idea that the increased fidelity of RNAPII around the stop codon is the result of reduced elongation rates.

**Figure 4.**
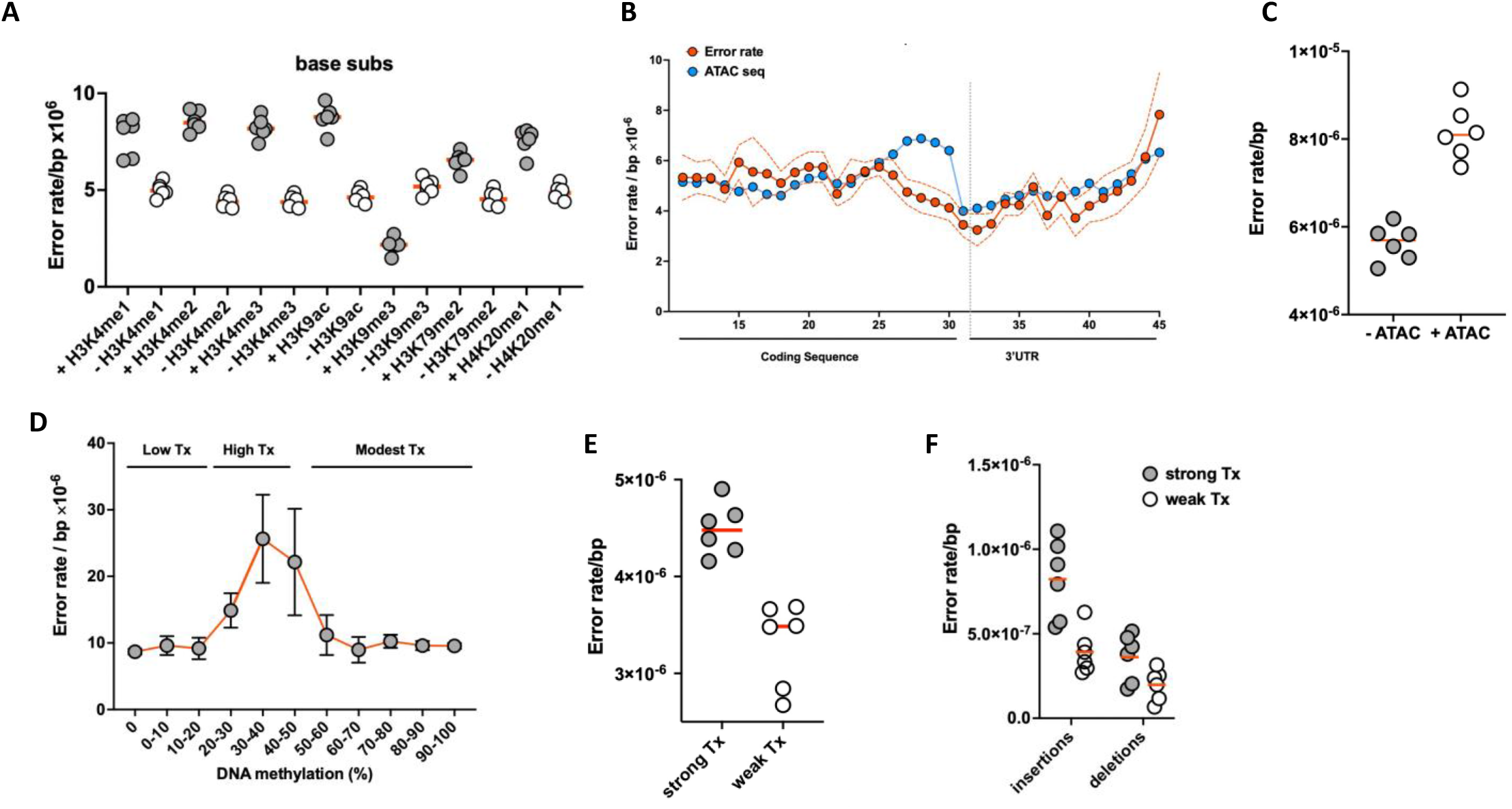
Epigenetic determinants of the error rate of transcription. **A**. Various histone modifications associated with active gene expression and RNAP elongation correlate with increased error rates, while markers that are associated with gene repression correlate with lower error rates. **B-C** Open sequences of DNA tend to display increased error rates. Please note that the ATAC-seq peaks displayed in panel B have been normalized to the error rate to better display their qualitative overlap. **D** Coding sequences that exhibit methylation levels associated with strong transcription display increased error rates. **E-F**. Genes that display strong transcription display higher error rates than those with weak transcription. In **4A, 4C, 4E** and **4F**, each datapoint represents one of the 6 replicates.

To further investigate the link between gene expression and fidelity, we examined how DNA accessibility affects the error rate of transcription. In support of the idea that increased expression compromises the fidelity of transcription, we found that ATAC-peaks (locations along the genome that are accessible to transposon integration) closely correlate with increased error rates along the length of a gene (**fig. 4B**), except for the reduced fidelity seen in 3’ end of the coding region. The more accessible the DNA is, the higher the error rate is for base substitutions, insertions and deletions (**fig. 4C**). This association is especially strong for specific types of errors like G→A substitutions, although other errors like U→C substitutions are anti-correlated with ATAC-peaks (**fig. S8**). More work will be needed to dissect the molecular basis for these observations. Another epigenetic mechanism associated with transcriptional activity is DNA methylation. Increased DNA methylation in the promoter, or 5’ UTR of a gene is usually associated with transcriptional repression. However, DNA methylation in the gene body has a parabolic relationship with gene expression, so that very little, or highly abundant methylation is associated with lower levels of transcription, while intermediate methylation is associated with active transcription^30^. Further supporting the idea that the strength of gene expression is directly coupled its fidelity, we found that the highest error rates occurred in genes with intermediate methylation levels in the CDS (**fig. 4D**). We then cemented this idea with a transcriptome-wide analysis of the ENCODE project dataset that showed that areas of strong transcription are associated with increased error rates for base substitutions, insertions and deletions (**fig. 4F-G**).

### BRCA1 controls the fidelity of transcription at R-loops

In addition to histones and epigenetic marks, gene bodies are also covered by various DNA-binding proteins. To determine if these proteins can affect the error rate of transcription, we monitored how the presence or absence of CTCF, MYC, Nanog, SIRT6, RAD21 and BRCA1 impact transcriptional mutagenesis. Although most of these proteins had little or no effect on transcriptional fidelity (**fig. S9**), we found that locations marked by BRCA1 displayed a significant increase in the error rate of transcription (**fig. 5A**). BRCA1 is best known as a DNA repair protein that plays a role in the suppression of breast cancer^31^. However, BRCA1 also binds to R-loops^32^, triple-stranded structures^33^ that are highly sensitive to DNA damage^34^. BRCA1 prevents these damaged structures from spreading across the genome and assists with local DNA repair processes if needed. Because DNA damage is a powerful source of transcription errors^15^, we hypothesized that damaged R-loops could be responsible for the increased error rates detected at BRCA1 peaks. To explore this hypothesis, we carefully examined the error spectrum and found that at BRCA1 peaks RNAPII displays a 5-fold increase in G→A errors and a 16-fold increase in C→A errors (**fig. 5B**). These errors are the two most common mistakes made by RNAPII on oxidatively damaged bases^15^. The most common consequence of oxidative damage is cytosine deamination, which creates uracil and uracil glycol lesions that mispair with adenine during transcription to induce G→A errors^35,36^. Similarly, oxidative damage causes 8-oxo-guanine lesions that mispair with adenine to induce C→A errors^8^. It was previously shown that by preventing damage-prone R-loops from spreading across the genome, BRCA1 limits mutagenesis in these structures. Our data now suggests that this function of BRCA1 also accomplishes a secondary goal, which is improving the fidelity of transcription.

**Figure 5.**
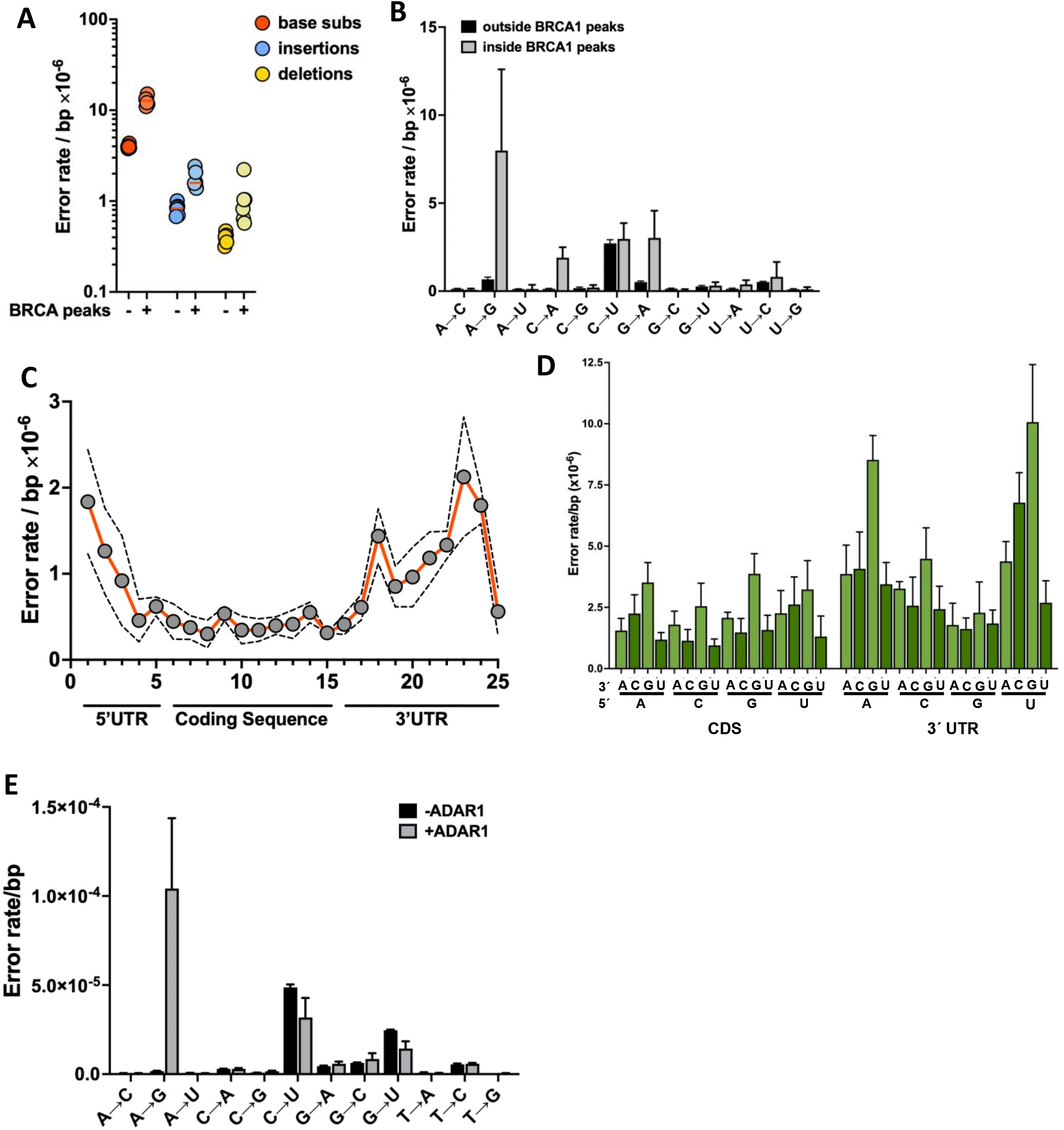
BRCA1 is associated with altered error rates and RNA editing. **A-B**. Genetic locations where BRCA1 is known to bind display elevated error rates and altered spectra. **C**. A→G “errors” incease in the 5’ and 3’ UTRs of genes, where A→I editing is known to occur. **D**. The genetic context of errors that replace adenine changes in the 3’ UTR towards the genetic context preferred by ADAR1. **E**. Expression of the human ADAR1 gene results in an elevated A→G error rate.

### Circle-sequencing allows for ultra-sensitive detection of off-target RNA editing

In addition to G→A and C→A errors, we observed an increase in A→G errors at BRCA1 peaks (**fig. 5B**). Interestingly, it was previously shown that R-loops at telomeres can be edited by ADAR1^37^, leading to A→I edits^38^ and a large body of evidence now supports the idea that these RNA editing enzymes tend to exhibit off-target effects. Because inosine pairs with cytosine, such off-target editing events on genomic R-loops could be reported as A→G errors by our circ-seq assay. In addition, it should be noted that A→I editing events are also common in the 3′ UTR of genes^39^. To test the idea that off-target A→I editing events are reported as A→G errors by our assay, we analyzed the location of A→G errors along gene bodies. We found that although this error is relatively rare in coding regions, it becomes increasingly prevalent in the 3′ UTR of human transcripts (**fig. 5C**). No other error displays a similar distribution (**fig. S10**). In addition, we found that the genetic context of the errors we detected on adenine bases in the 3′ UTR shifts towards the genetic context that is preferred by ADAR1, which prefers an adenine or uracil as it’s 5′ nearest neighbor and a guanine on it’s 3′ side^40^ (**fig. 5D**). Taken together, these observations suggest that circle-sequencing is detecting editing events that are performed by ADAR enzymes. To explore this hypothesis further, we expressed the human ADAR1 gene in the budding yeast *S. cerevisiae* and detected a clear increase in A→G errors (**fig. 5E**). This observation unambiguously demonstrates that circle sequencing can detect RNA editing events across the transcriptome, lending credence to the idea that the A→G errors detected at BRCA1 peaks are indeed off-target editing events. It should be noted though, that all the edits detected this way are likely to be off-target editing events. The safeguards present in our bio-informatic pipeline to prevent editing events from being reported as transcription errors ensure that any site that is edited >0.5% is automatically discarded. To be included into our analysis, each base pair must be sequenced >200 times, and if an edit occurs in more than 0.5% of these reads, it is automatically discarded. It is probably most prudent then, to call these errors transcript errors, as opposed to transcription errors. Intriguingly though, editing errors are now recognized as a new type of “biological error” that is receiving increasing attention due to its potential physiological consequences on protein aggregation and the generation of unique, immunogenic epitopes. Importantly, transcription errors have similar effects on cells. Our observations now suggest that circle-sequencing could provide a new tool to catalogue and analyze off-target editing across the transcriptome, so that their impact on human biology can be evaluated. Those experiments are especially important in the context of the genetically engineered RNA editors that are currently being tested for therapeutic purposes in humans. Circle-sequencing could provide an essential quality control mechanism to ensure that these editors result in as few off-target effect as possible.

## DISCUSSION

The accuracy of DNA replication, transcription and translation form the foundation of life itself. Together, these processes ensure the faithful inheritance and expression of our genetic code. It is essential therefore, that we understand the molecular mechanisms that control the fidelity of these processes. Surprisingly though, relatively little is known about the fidelity of transcription, leaving a key component of life largely unexplored. To fill this gap in our knowledge we measured the fidelity of transcription in human embryonic stem cells. These measurements demonstrate that the error rate of transcription is remarkably variable across the transcriptome. The error rate does not only differ between polymerases, but also between classes of genes and even with specific regions of these genes. One surprising conclusion then, is that human cells seem to prioritize the fidelity of some transcripts over others. Our analyses indicate that they are likely to do so by two separate mechanisms: by assembling RNAPII with or without fidelity factors, or by altering the speed of RNA synthesis. First, we found that the error spectrum of ribozyme RNAs perfectly mimics the error spectrum of *S. cerevisiae* cells that lack the fidelity factor TFIIS and *C. elegans* worms that lack the TFIIS homologue T24H10.1. Human cells carry 3 homologues of TFIIS and T24H10.1, labelled TCEA1, 2 and 3. Based on our data, we predict that these factors are not part of the RNAPII holoenzyme that transcribes ribozymes. In addition, we found that lncRNAs display an increased error rate for all possible base substitutions, insertions and deletions, suggesting that the inherent fidelity of RNAPII itself has changed. Our observations regarding epigenetic markers, the error rate near stop codons and the genetic context of error rates, suggests that genes might be able to modify their error rate by altering the speed of RNAPII elongation. This phenomenon seems to be part of a broader narrative that suggests that the strength of transcription is directly linked to fidelity. Intriguingly, genes that are rarely transcribed tend to display lower error rates than genes that are highly transcribed. The improved fidelity of rarely transcribed genes could have a clear advantage for human cells. If few transcripts are made of a gene, an error in one of those transcripts could have an outsized impact on its function. Accordingly, the remarkable correlation between the strength of transcription and its fidelity could be the result of a co-evolutionary process. Our data also suggests that BRCA1 may have an additional, uncharacterized function in controlling the fidelity of transcription. BRCA1 is known to bind to R-loops^32^, triple-stranded structures that are prone to the accumulation of DNA damage^33^. BRCA1 prevents these R-loops from spreading^32^ and assists with DNA repair processes if needed. Accordingly, BRCA1 limits DNA damage from accumulating across the genome and prevents mutations from arising. It now seems that these functions have a secondary purpose as well: to improve the fidelity of transcription. In this context, it is important to note that DNA damage-induced transcription errors were previously shown to activate oncogenic pathways in human cells^8^. Accordingly, our observations suggest that one additional mechanism by which BRCA1 suppresses human cancers is the prevention of transcriptional mutagenesis. Another consequence of transcription errors is protein aggregation^1,7,9^. It was previously shown that slippage events on dinucleotide tracts in the *APP* and *UBB* gene result in toxic peptides that are part of the amyloid plaques that characterize Alzheimer’s disease. In addition, it was shown that the UBB^+1^ peptide is a potent inhibitor of the proteasome ubiquitin complex^3^ and can also be found in protein aggregates from patients with Guam parkinsonism–dementia complex, tauopathies and polyglutamine repeat disorders^1,4-6^, suggesting that errors in the *UBB* transcript could contribute to other protein misfolding diseases as well. Here, we build on these findings by demonstrating that mono- and di-nucleotide tracts, like those present in the APP and UBB transcripts are especially dangerous because they display greatly increased error rates. In addition, we found that these types of slippage events occur most frequently in cells that are directly implicated in the progression of Alzheimer’s disease. These two observations suggest that cells involved in Alzheimer’s disease experience a constant stream of toxic peptides that promote the disease, strongly supporting the idea that transcription errors play a role in disease progression. It will be interesting to examine this possibility in future research.

## Funding

This research was supported by grants from the US Department of Army, MURI award W911NF-14-1-0411 (M.L.), National Institutes of Health, R35-GM122566-01 (M.L), the National Science Foundation, DBI-2119963 (M.L.), and the National Institute on Aging, R01AG054641 NIA (M.L, J-F.G and M.V.).

**Figure S1.**
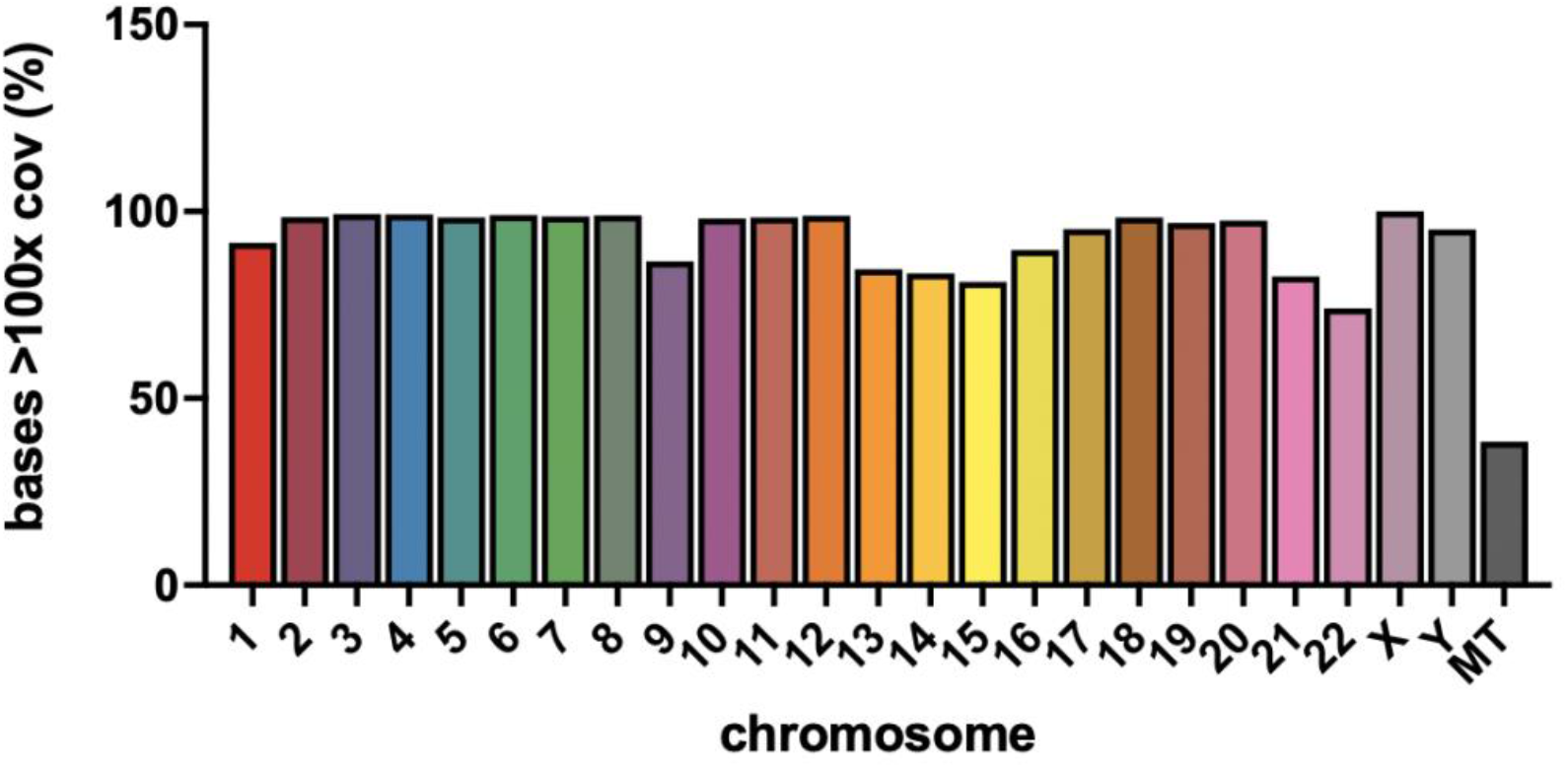
DNA coverage of H1 hESCs. This graph depicts the percentage of bases of each chromosome for which we have >100x coverage.

**Figure S2.**
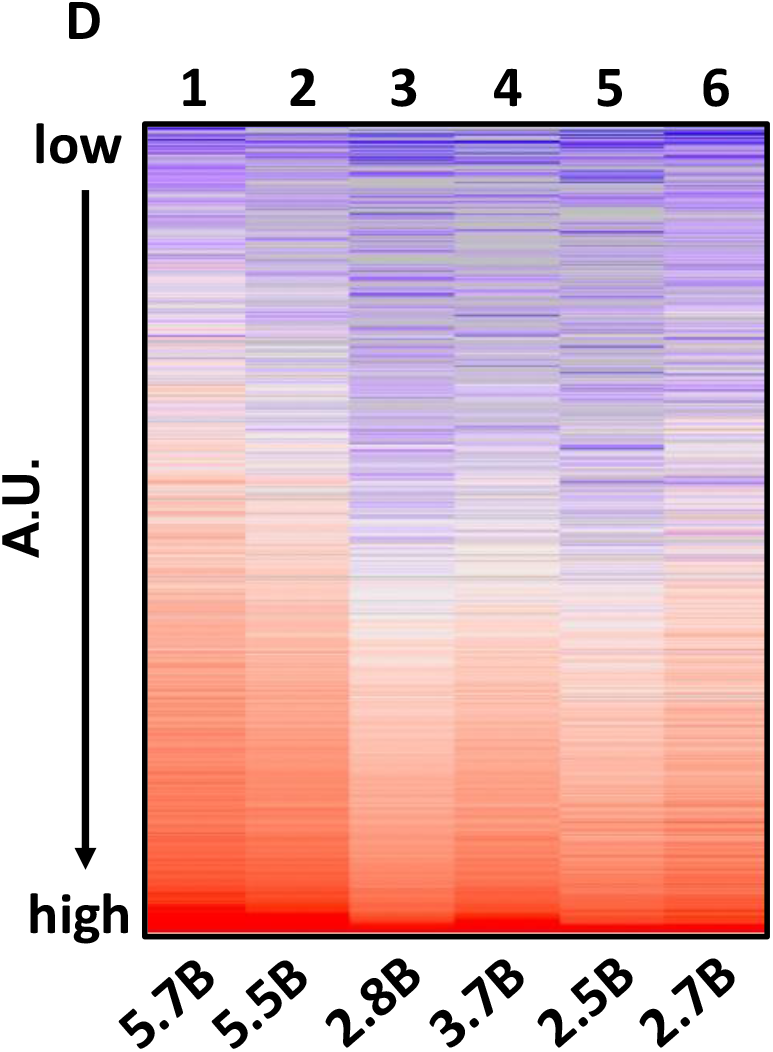
Transcriptome profile of the 6 H1 hESC replicates sequenced for this study. Transcriptome profiles of each replicate (numbering on top) were lined up so that each line corresponds to the same gene in each dataset. Expression levels are depicted in arbitrary units, where blue coding equals low expression and red coding equals high expression. All samples were normalized by reads per kilobase of transcripts per million mapped reads. The number of bases sequenced from each replicate are depicted below the graph. In total 9,437 genes were detected at least once over all six samples.

**Figure S3.**
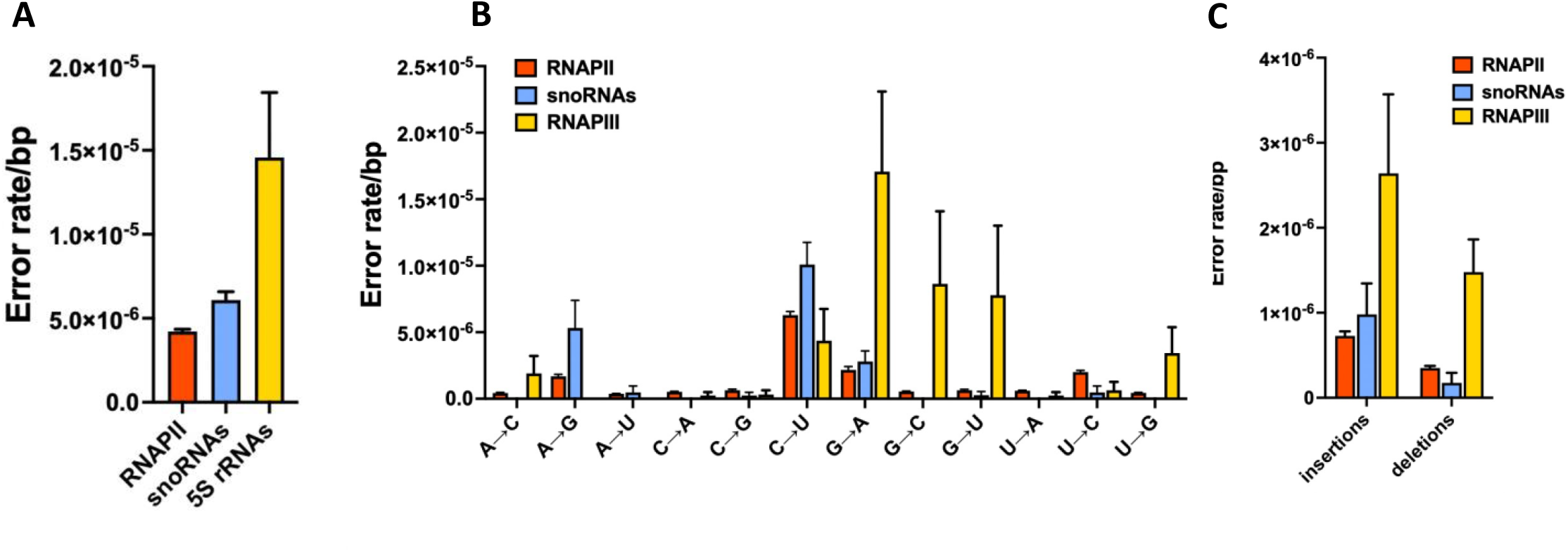
snoRNAs display a different error spectrum compared to RNAPIII. **A**. The error rate of snoRNAs is significantly than other genes known to be transcribed by RNAPIII. **B-C**. The error spectrum of snoRNAs is significantly different from the spectrum of other genes known to be transcribed by RNAPIII and resembles the error rate and spectrum of RNAPI and II better.

**Figure S4.**
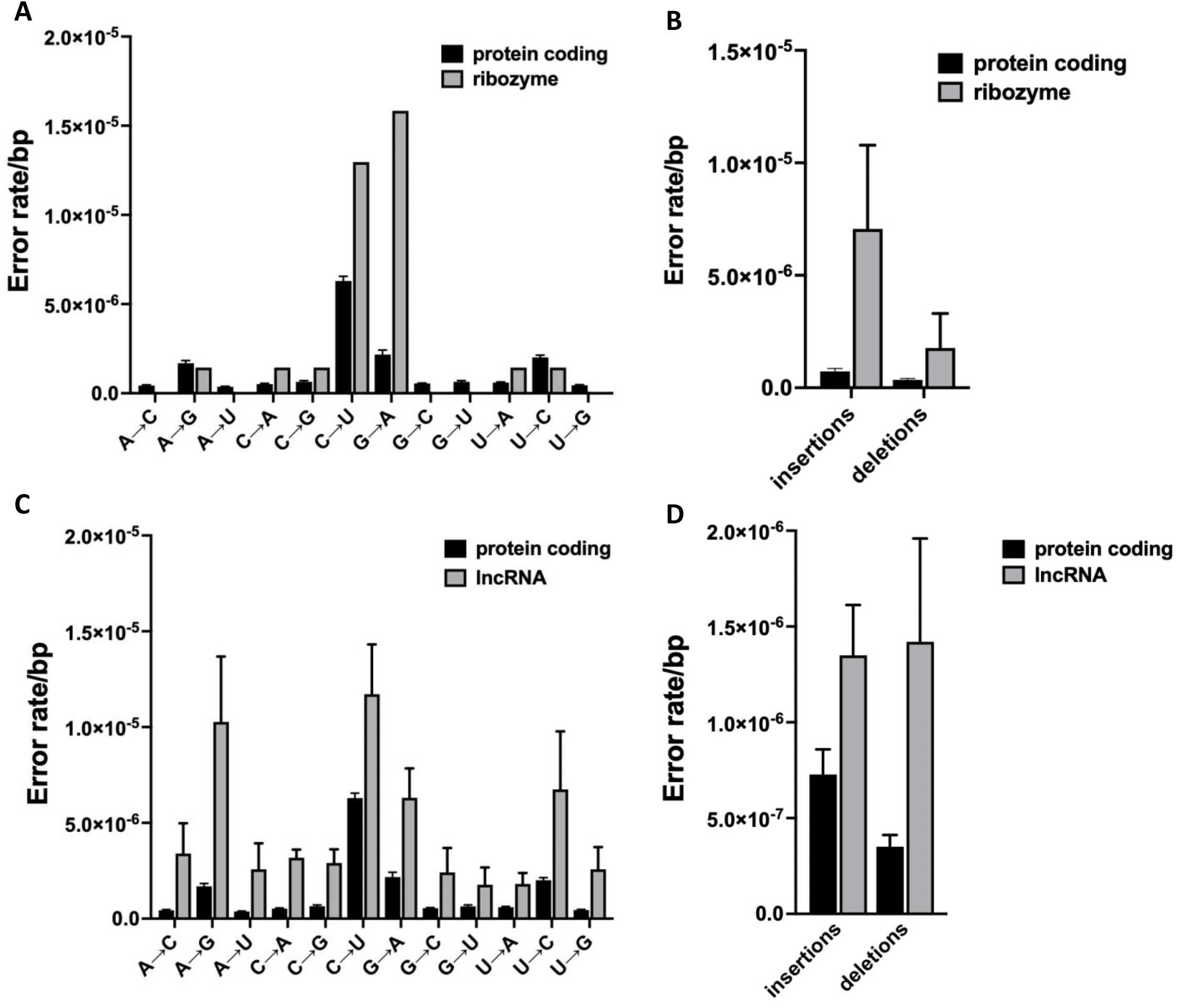
Increased error rates in ribozymes and lncRNAs. **A-B**. The increased error rate of ribozymes is primarily caused by an increased number of G→A errors and insertions and deletions, strongly resembling the transcription of mRNAs by RNAPII that lack a fidelity factor. Please note that in panel **A** the base substitutions detected in ribozymes in all replicates have been combined due to the relatively low coverage of ribozymes across the transcriptome. **C-D**. In lncRNAs every possible class of errors is elevated.

**Figure S5.**
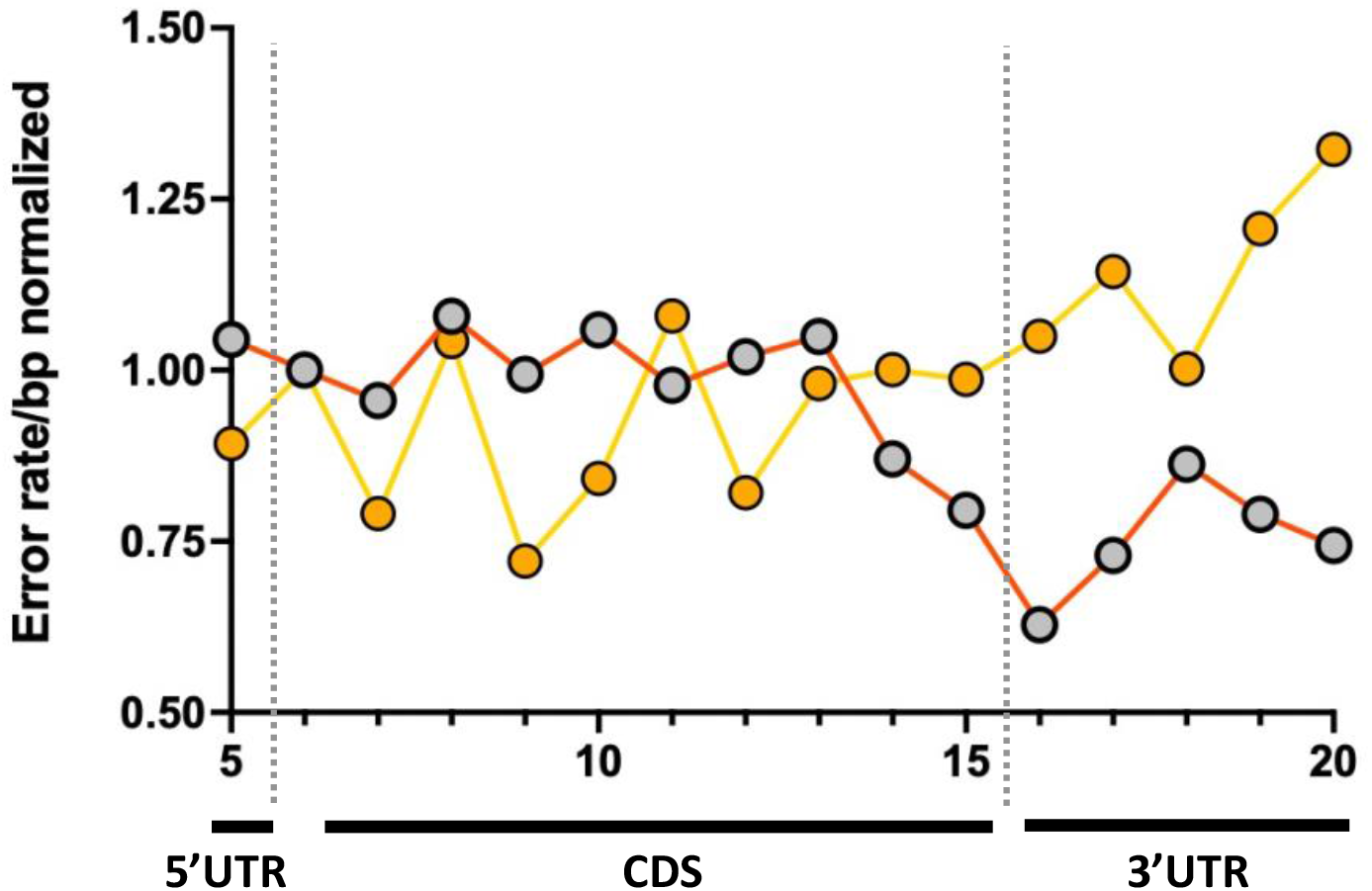
Deletions along the length of genes. In contrast to base substitutions, deletions increase near the stop codon of protein coding genes. Please note that the deletion and base substitution rate have been normalized to each other to better display their qualitative differences.

**Figure S6.**
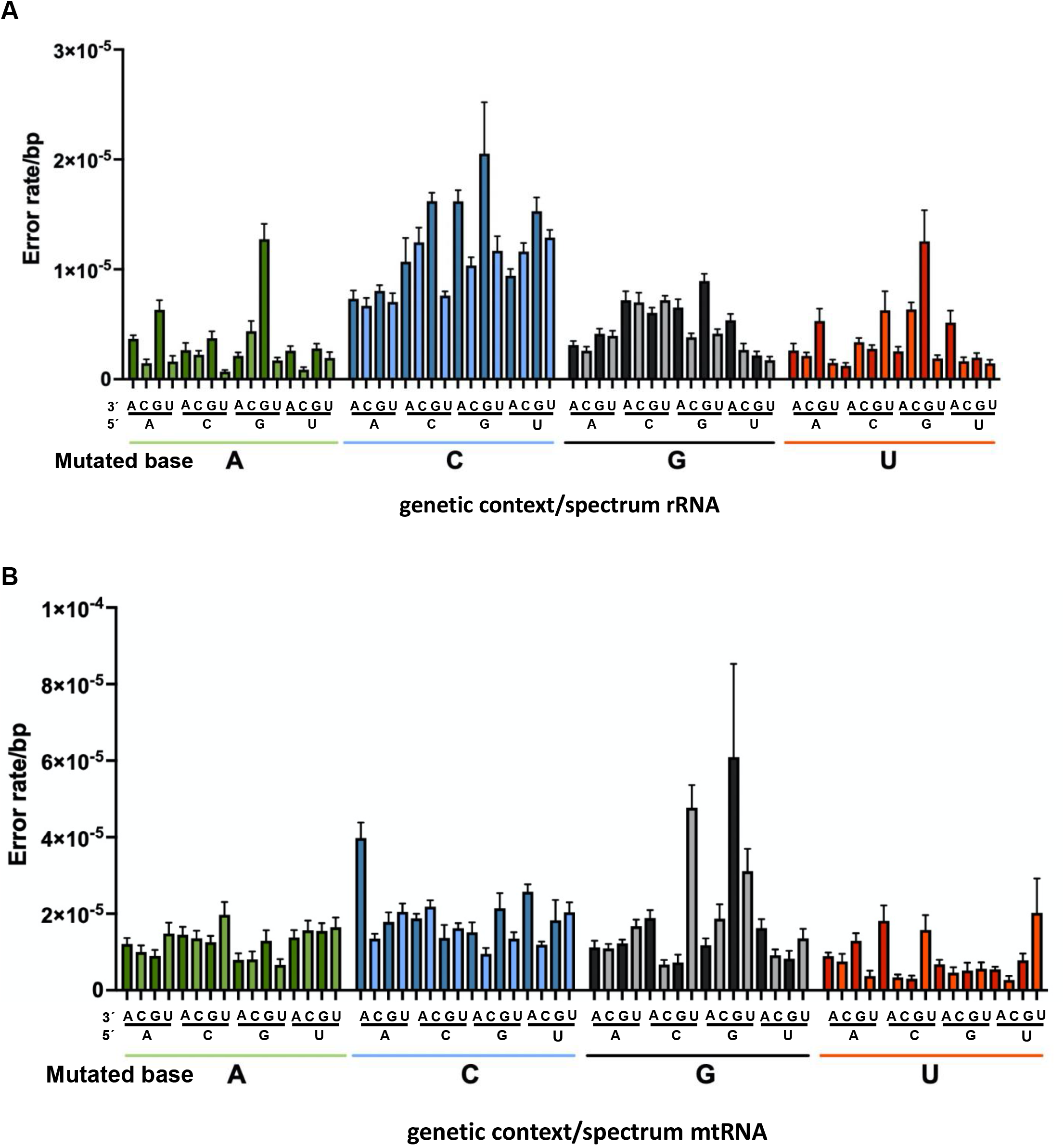
Error spectrum as a function of genetic context. **A**. Error spectrum of rRNA, synthesized by RNAPI. Similar to RNAPII, the error rate tends to be higher if the next RNA base that needs to be inserted is a purine. Or in other words, if the next DNA base that needs to be transcribed is a pyrimidine. **B**. Error spectrum of mtRNA, synthesized by the mitochondrial RNA polymerase. In contrast to

**Figure S7.**
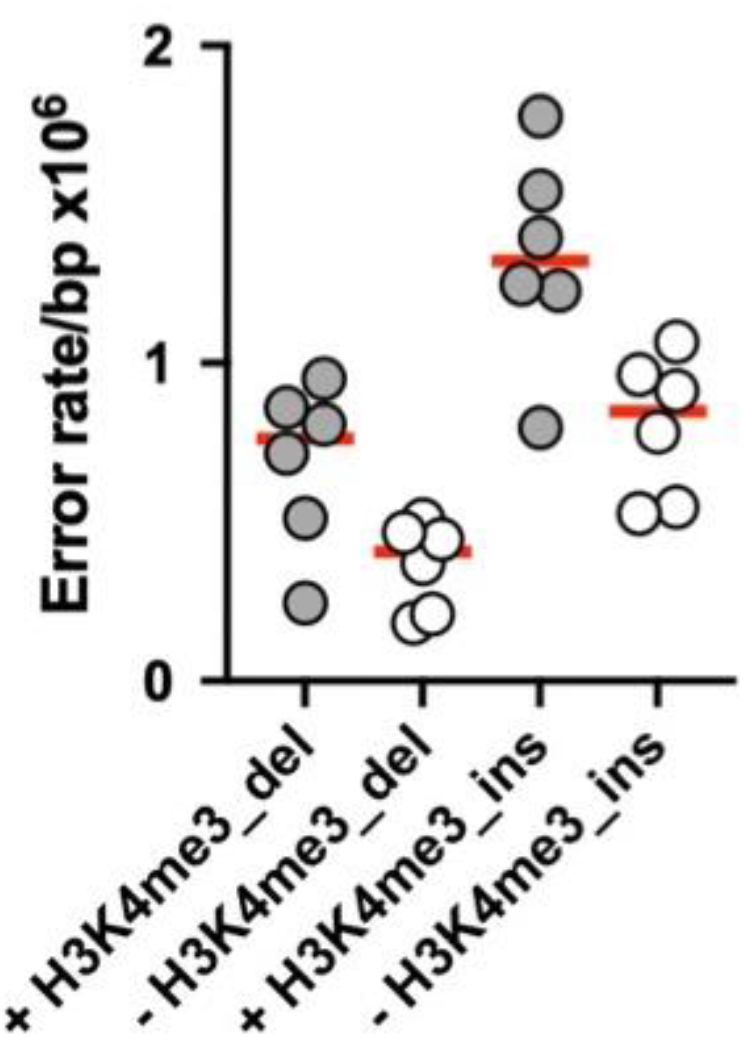
Histone modifications and the error rate of transcription. The active expression marker H3K4me3 is correlated with an elevated rate of insertions and deletions.

**Figure S8.**
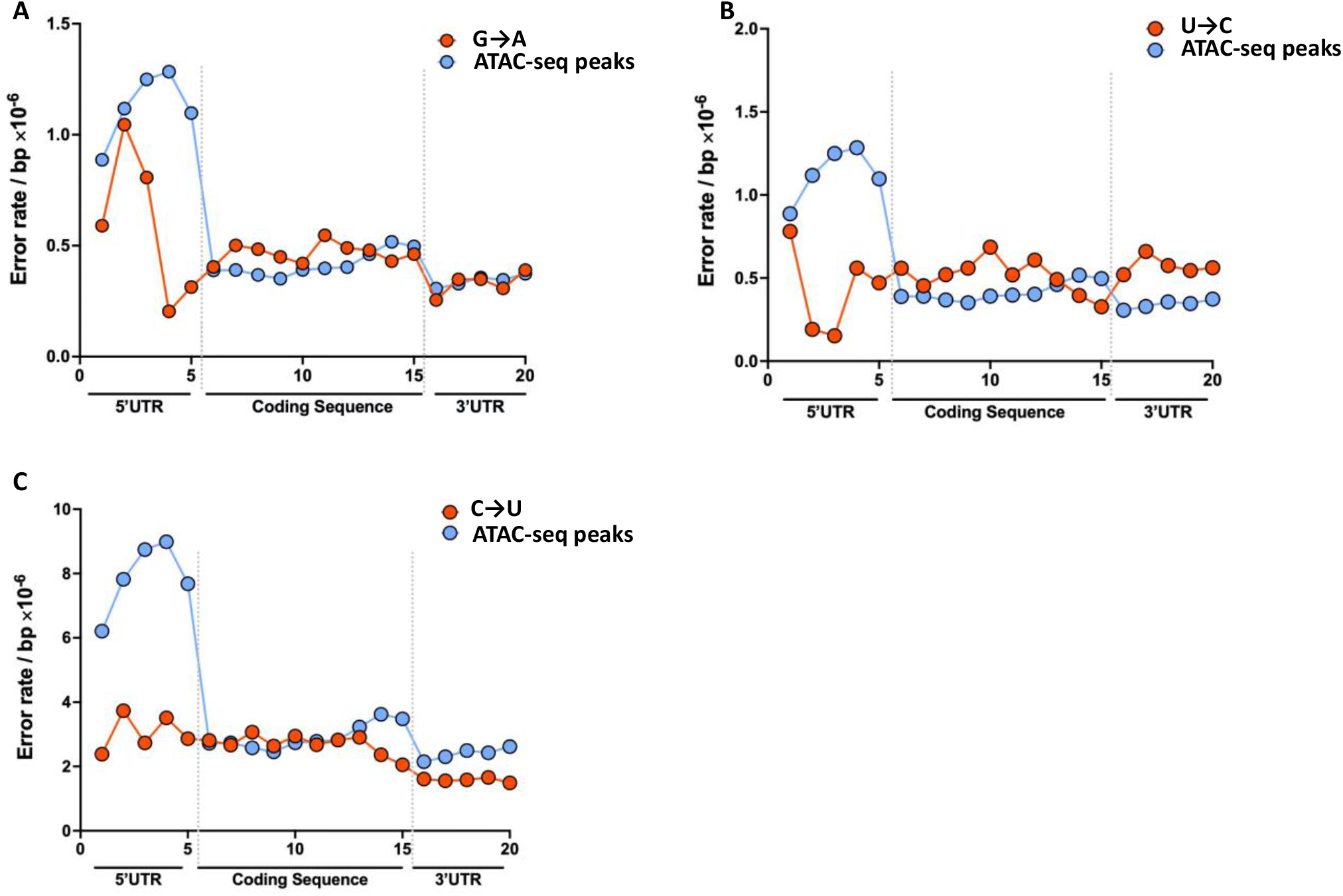
The correlation of errors with ATAC-seq peaks. **A**. G→A errors are closely correlated with the openness of DNA. **B**. U→C errors are anti-correlated with the openness of DNA. C→U errors neither correlate or are anti-correlated with open DNA. Please note that the ATAC-seq peaks displayed in all panels have been normalized to the error rate to better display their qualitative overlap (or lack thereof).

**Figure S9.**
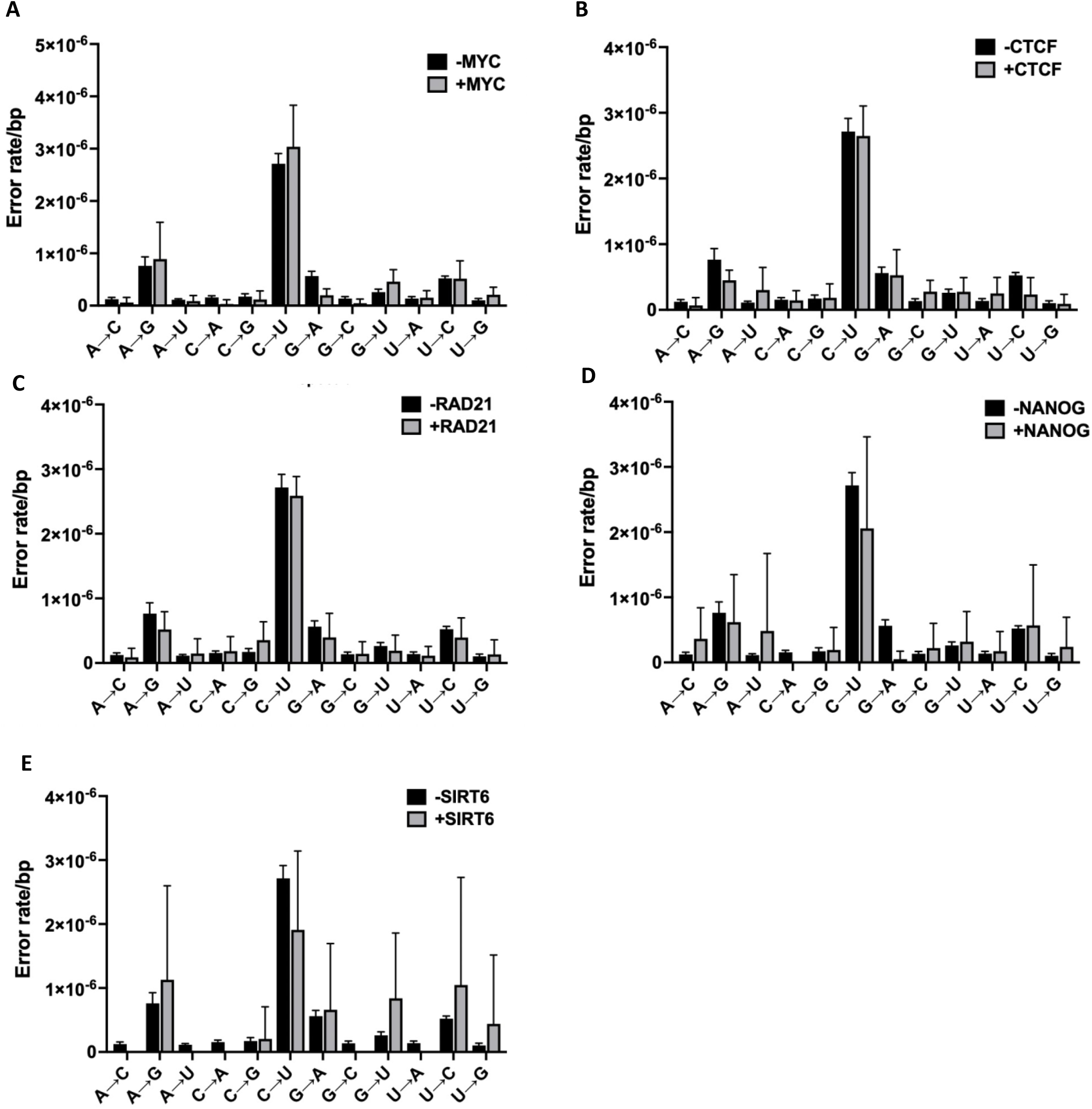
DNA binding proteins and the error rate of transcription. **A-E**. The presence of MYC, CTCF, RAD21, NANOG and SIRT6 has no impact on the error rate of transcription.

**Figure S10.**
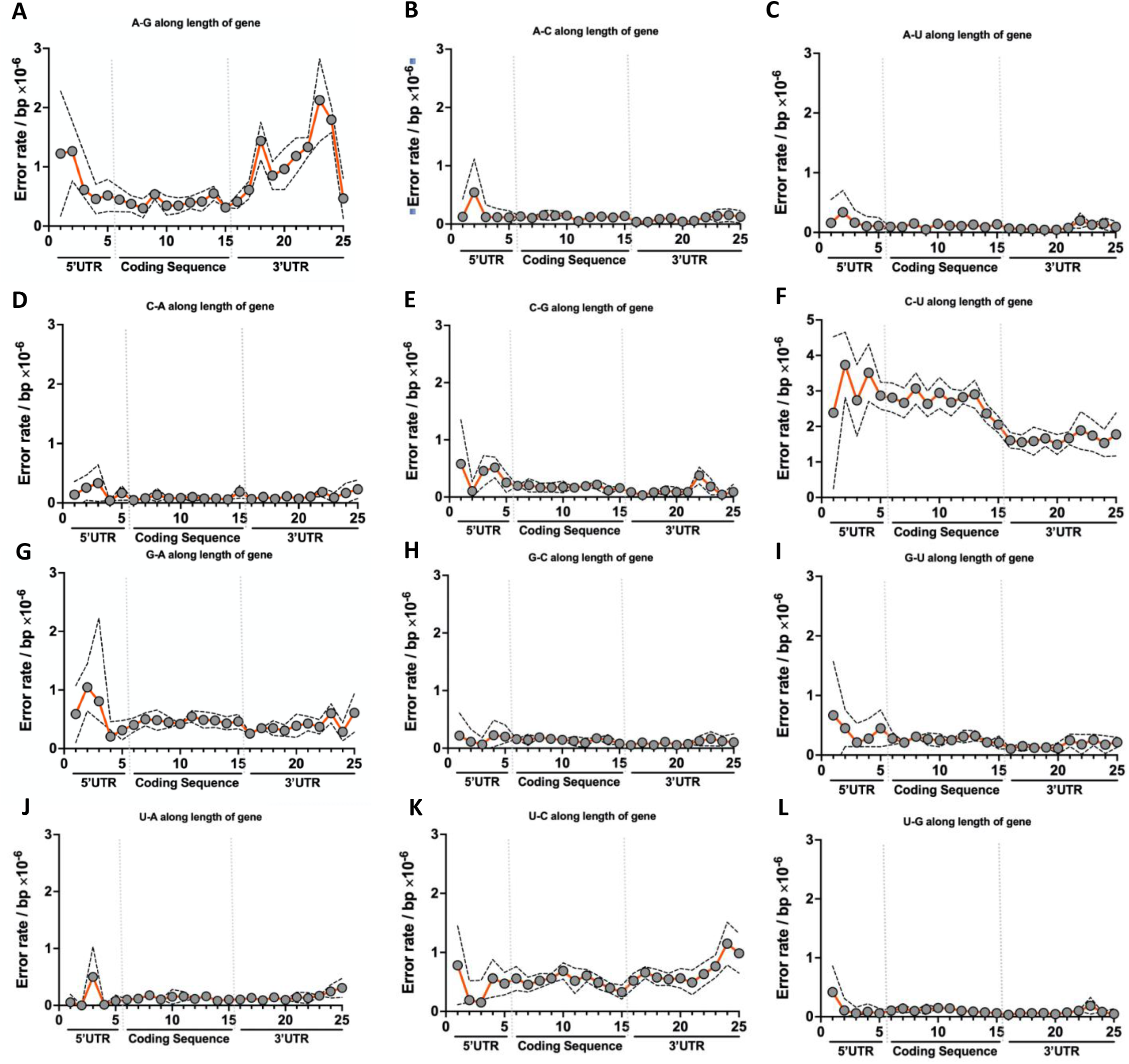
The correlation of all possible base substitutions along the length of genes. **A-L**. Only A→G errors are increasingly (and significantly) present in both the 5’ and 3’ UTR of genes.

## METHODS

### H1 hESC Cell culture

H1 hESCs were purchased from WiCell in Wisconsin (WA01) and cultured in TeSR medium in Matrigel coated 10cm plates. Cells were grown at 5% O^2^ tension to better mimic the conditions inside the human body and reduce oxidative damage as a result of normoxic conditions. To passage cells and prior to collection of RNA and DNA, cells were gently treated with 2 μg/ml Dispase mixed with DMEM/F12, washed with PBS and scraped off the plate using a glass pipette. DNA and RNA were then isolated with standard phenol chloroform and Trizol methods.

### Library Construction and Sequencing

Library preparation 1100ng of enriched mRNA was fragmented with the NEBNext RNase III RNA Fragmentation Module (E6146S) for 25 minutes at 37°C. RNA fragments were then purified with an Oligo Clean & Concentrator kit (D4061) by Zymo Research according to the manufacturer’s recommendations, except that the columns were washed twice instead of once. The fragmented RNA was then circularized with RNA ligase 1 in 20 μl reactions (NEB, M0204S) for 2 hours at 25°C after which the circularized RNA was purified with the Oligo Clean & Concentrator kit (D4061) by Zymo Research. The circular RNA templates were then reverse transcribed in a rolling-circle reaction by first incubating the RNA with for 10 minutes at 25°C to allow the random hexamers used for priming to bind to the templates. Then, the reaction was shifted to 42°C for 20 minutes to allow for primer extension and cDNA synthesis. Second strand synthesis and the remaining steps for library preparation were then performed with the NEBNext Ultra RNA Library Prep Kit for Illumina (E7530L) and the NEBNext Multiplex Oligos for Illumina (E7335S, E7500S) according to the manufacturer’s protocols. Briefly, cDNA templates were purified with the Oligo Clean & Concentrator kit (D4061) by Zymo Research and incubated with the second strand synthesis kit from NEB (E6111S). Double-stranded DNA was then entered into the end-repair module of RNA Library Prep Kit for Illumina from NEB, and size selected for 500-700 bp inserts using AMPure XP beads. These molecules were then amplified with Q5 PCR enzyme using 11 cycles of PCR, using a two-step protocol with 65°C primer annealing and extension and 95°C melting steps.sequencing data was converted to industry standard Fastq files using BCL2FASTQv1.8.4.

### Error Identification

We have developed a robust bioinformatics pipeline to analyze circ-seq datasets and identify transcription errors with high sensitivity^9,41^. First, tandem repeats are identified within each read (minimum repeat size: 30nt, minimum identity between repeats: 90%), and a consensus sequence of the repeat unit is built. Next, the position that corresponds to the 5′ end of the RNA template is identified (the RT reaction is randomly primed, so cDNA copies can “start” anywhere on the template) by searching for the longest continuous mapping region. The consensus sequence is then reorganized to start from the 5′ end of the original RNA fragment, mapped against the genome with tophat (version 2.1.0 with bowtie 2.1.0) and all non-perfect hits go through a refining algorithm to search for the location of the 5′ end before being mapped again. Finally, every mapped nucleotide is inspected and must pass **5** checks to be retained: **1)** it must be part of at least 3 repeats generated from the original RNA template; **2)** all repeats must make the same base call; **3)** the sum of all qualities scores of this base must be >100; **4)** it must be >2 nucleotides away from both ends of the consensus sequence; **5)** each base must be covered by ≥200 reads with <1% of these reads supporting a base call different from the reference genome. This final step filters out polymorphic sites and potential RNA-editing events. For example, if a base call is different from the reference genome, but is present in 100 out of 200 reads, it is not labeled as an error but as a heterozygous mutation. A similar rationale applies to low-level mutations and RNA editing events. Each read containing ≥1 mismatch is filtered through a second refining and mapping algorithm to ensure that errors in calling the position of the 5′ end cannot contribute to false positives. The error rate is then calculated as the number of mismatches divided by the total number of bases that passed all quality thresholds.

### Bio-informatic analysis

To determine the location of transcription errors along the length of genes, we first mapped each error to a distinct coordinate along the length of the human genome, using the 38^th^ assembly of the human genome as our guide (GRCh38). We then generated the circle plots in figure 1 with the software “circa”, which was downloaded from https://omgenomics.com/circa/. Then, we cross-referenced these coordinates with the gene and transcript annotations provided by the 100^th^ version of Ensembl. Once we knew the transcripts that were affected by each error, we determined the exact position of each error within these transcripts with the GenomicFeatures package in R. Finally, we separated the transcripts into their 5’UTR, coding region and 3’UTR, and divided these regions into bins. We then used the number of errors that were assigned to each bin to compute the error rate along the length of the entire transcript.

To determine the role of histone modifications and transcript factors on the error rate of transcription, we first downloaded the relevant histone modification and transcript factor ChIP-Seq data from the ENCODE (https://www.encodeproject.org), NIH Roadmap Epigenomics Mapping Consortium (http://www.roadmapepigenomics.org/data/), and Cistrome databases (http://cistrome.org/). When multiple datasets were available for the same parameter, we compared them to each other and identified overlapping peaks that were shared between datasets. We then used these overlapping peak calls to separate transcription events as within or outside of the peak regions. We then calculated the error rates as above for both peak and non-peak regions.

To determine the impact of DNA methylation on the error rate of transcription, we downloaded the processed, whole genome bisulfite sequencing (WGBS) dataset in H1 hESCs from: https://egg2.wustl.edu/roadmap/web_portal/processed_data.html#MethylData. Because this dataset was mapped onto a previous version of the human genome assembly (hg19) we first lifted it over to the current assembly (hg38). Then, we correlated the frequency of methylation at CpG sites with the error rate of transcription. To do so, we divided the methylation frequency of CpG sites into 11 bins, where bin 1 contains all sites that display no methylation, and bin 2-10 contains CpG sites that are methylated in 10% increments, so that bin 2 = 0-10% mythlation, bin 2 10-20% methylation and son on. We then calculated the error rates for each of these bins as described above.

To determine how strong or weak transcription affects the error rate, we downloaded the core 15-state model data is downloaded from: https://egg2.wustl.edu/roadmap/web_portal/chr_state_learning.html#core_15state. We then assigned the transcription errors and the number of times each base was covered to each core state and calculated the error frequency as above.

### Staining of mouse tissues

Mice were kept on a 12-hour light/dark cycle with food and water available ad libitum. Only male mice were used for experiments. Experimental mice were euthanized by CO_2_ asphyxiation and cervical dislocation, and tissues were immediately collected and fixed in 10% neutral buffered formalin (15740-01, EMS, Hatfield, PA, USA) for 24h at RT. Formalin-fixed tissues were rinsed with PBS and cryoprotected with 30% sucrose in PBS at 4°C. Next, tissues were embedded with Tissue-Tek OCT compound (4583, Sakura Finetechnical, Tokyo, Japan) and rapidly frozen in chilled isopentane/2-methylbutane. Frozen OCT-embedded tissues were sectioned on a cryostat (Leica CM1860, Leica Biosystems, Wetzlar, Germany) into 10 μm-thick sections and collected on Superfrost Plus microscope slides (12-550-15, Fisher Scientific). For immunofluorescence, sections were incubated with rabbit anti-GFP primary antibodies (1:500, 600-401-215, Rockland, USA) overnight at 4°C. Next, sections were rinsed with PBS and incubated with Alexa Fluor 488 goat anti-rabbit IgG secondary antibodies (1:400, A11034, Invitrogen) for 1h at RT. Sections were subsequently rinsed with PBS and mounted with Vectashield mounting media with DAPI (H-1200, Vector laboratories, USA). Slides were imaged on an ECHO microscope or a Zeiss LSM780 confocal. Tissue sections from Sox2Cre x YFP^neo^ mice were used as positive controls. All animal experiments were performed in compliance with the guidelines set out by the Institutional Animal Care and Use Committee.

## Notes

### Competing Interest Statement

The authors have declared no competing interest.

